# Dynamic changes in oligomeric complexes of UPR sensors induced by misfolded proteins in the ER

**DOI:** 10.1101/189605

**Authors:** Arunkumar Sundaram, Suhila Appathurai, Malaiyalam Mariappan

## Abstract

The endoplasmic reticulum (ER) localized unfolded protein response (UPR) sensors, IRE1α, PERK, and ATF6α, are activated upon accumulation of misfolded proteins caused by ER stress. It is debated whether these UPR sensors are activated either by the release of their negative regulator BiP chaperone or directly binding to misfolded proteins during ER stress. Here we simultaneously examined oligomerization and activation of all three endogenous UPR sensors. We found that UPR sensors existed as preformed oligomers even in unstressed cells, which shifted to large oligomers for PERK and small oligomers for ATF6α, but little changed for IRE1α upon ER stress. Neither depletion nor overexpression of BiP had significant effects on oligomeric complexes of UPR sensors both in unstressed and stressed cells. Thus, our results find less evidence for the BiP-mediated activation of UPR sensors in mammalian cells and support that misfolded proteins bind and activate the preformed oligomers of UPR sensors.

## Introduction

The endoplasmic reticulum is the major organelle for the synthesis of secretory and membrane proteins. These proteins enter the ER through the Sec61 translocon channel and mature with the help of cascade of chaperones, folding enzymes and post-translocation modifications (Rapoport, 2007; van Anken 2005). The proteins that fail to achieve their native state are recognized and eliminated by the ER associated degradation (ERAD) pathways (Brodsky, 2012; Christianson and Ye, 2012). Thus, only folded proteins are packaged into vesicles for their transport to the Golgi apparatus. However, environmental stress, nutrient protein overload, or expression of mutant proteins overwhelms ERAD machinery, thus leading to accumulation of misfolded proteins in the ER. The excess of misfolded proteins activates the conserved unfolded protein response (UPR) pathway, which transmits the information of the folding status of the ER to the cytosol and nucleus (Walter and Ron, 2011). This leads to activation of transcriptional and translational programs to increase the ER protein folding capacity by upregulating chaperones folding enzymes, and ERAD machinery (Lee et al., 2003; Shoulders et al., 2013). In case of failure to attenuate the UPR due to prolonged stress, the cells commit suicide by initiating apoptotic pathways. The dysfunction or overactive UPR signaling has been implicated in numerous human diseases including type 2 diabetes, neurodegenerative diseases, and cancer (Han et al., 2009; Hetz, 2012; Wang and Kaufman, 2016).

In metazoans, three UPR sensors, IRE1α, PERK and ATF6α are known to detect the accumulation of misfolded proteins in the lumen and transmit the signal to the cytosol (Walter and Ron, 2011). IRE1α is a transmembrane kinase/endonuclease (RNase) that, upon ER stress, initiates the unconventional splicing of XBP1 mRNA (Cox et al., 1993; Mori et al., 1993; Yoshida et al., 2001; Calfon et al., 2002). The spliced XBP1 mRNA encodes an active transcription factor that upregulates genes such as chaperones and ER degradation machinery to improve the ER protein folding capacity (Lee et al., 2003; Shoulders et al., 2013). In addition, IRE1α can also reduce protein synthesis load at the ER through promiscuously ER-localized mRNAs encoding membrane and secretory proteins, a process known as IRE1α-dependent mRNA decay (RIDD) (Hollien and Weissman, 2006; Hollien et al., 2009). PERK is a transmembrane kinase, and its luminal domain shares a limited homology (~12%) to the luminal domain of IRE1α (Zhou et al., 2006). Upon ER stress, PERK phosphorylates eukaryotic translation initiation factor to shut down the overall protein synthesis, thus counteracting protein overload at the ER (Harding et al., 1999; Sood et al., 2000). However, some mRNAs that have small open reading frames in their 5’untranslated regions are translated by phosphorylated eIF2α, thereby production of transcription factors such as ATF4 (Ameri and Harris, 2008). ATF6α is an ER-localized transmembrane transcription factor (Haze et al., 1999). During ER stress conditions, ATF6α transported to the Golgi apparatus, where its cytoplasmic domain is released from membrane domain by S1P and S2P-mediated proteolysis (Ye et al., 2000; Nadanaka et al. 2007; Shindler and Schekman, 2009). The cleaved ATF6α moves to the nucleus and drives transcription of genes encoding chaperones and ERAD machinery for restoring ER homeostasis (Lee et al., 2003; Shoulders et al., 2013).

While there is tremendous progress has been made in understanding the biology of UPR effectors, the mechanism of UPR sensors activation remains incompletely understood. There are two major models have been actively debated for the activation of UPR sensors (Walter and Ron 2011; Snapp, 2012). The first model is similar to other stress sensing pathways such as the heat shock response that is strongly regulated by the binding and availability of a chaperone (Arsene et al., 2000; Anckar et al., 2011). Accordingly, BiP binds with monomers of IRE1α and PERK, thus preventing oligomerization and activation in unstressed cells. During ER stress, BiP is sequestered by misfolded proteins, thus allowing IRE1α and PERK to freely diffuse, oligomerize and become activated. This model is supported by the evidence that IRE1α and PERK associate with BiP in unstressed cells and that the association is disrupted in the presence of ER stress (Bertolotti et al., 2000; Okamura et al., 2000; Oikawa et al., 2009; Carrara et al., 2015). The activation of ATF6α appears to be slightly different from other two sensors since it seems to form oligomers under unstressed conditions but associated with BiP (Nadanaka et al. 2007; Gallagher et al., 2016; Shen et al., 2002). Upon ER stress, ATF6α moves from the ER to the Golgi, which appears to correlate with the release of BiP from ATF6α (Shen et al., 2002).

In the second model, unfolded proteins may directly bind to the luminal sensor domains of UPR sensors with concomitant release of BiP from the luminal domains. This binding may drive oligomerization change and activation of UPR sensors (Walter and Ron, 2011). The first evidence supporting this model came from crystal structures of yeast Ire1p luminal domain, which resembles the peptide-binding groove of MHC-I (Credle et al., 2005). Based on this, the Peter Walter group proposed the peptide-binding hypothesis. This idea is corroborated by the detection of interaction between misfolded proteins and Ire1p (Kimata et al., 2007; Gardener et al., 2011). However, there are studies challenge the peptide-binding model. First, the human IRE1α luminal domain structure exhibits a narrow peptide-binding groove of MHC-1, which may not accommodate misfolded proteins, although a recent study suggests a mutation in the groove of MHC-1 seems to interfere with the detection of misfolded proteins in the ER lumen (Kono et al., 2017). Unlike yeast Ire1p, human IRE1α does not seem to interact with misfolded proteins (Oikawa et al., 2012). Second, the fact that monomeric form of IRE1α cannot bind to unfolded peptides in vitro raises the question of how monomers of IRE1α can efficiently bind to misfolded proteins in cells during ER stress conditions (Gardner and Walter, 2011).

It has been challenging to determine which of these models is occurring in mammalian cells. One of the key requirements to test these different models is to monitor the endogenous oligomeric complexes of all three UPR sensors under homeostatic and ER stress conditions in cells. Although size fractionation assays to probe the oligomerization of UPR sensors were successful, they were laborious to test different time points of stress since it involves examining several fractionated samples. This approach is further complicated by the fact that all three UPR sensors are relatively low abundant proteins in cells. Thus, it is not feasible to detect these proteins in diluted size-fractionated samples. An imaging-based approach that monitors ER stress dependent higher order oligomers (or clusters) proves to be useful for probing IRE1α activation in both in yeast and human cells. However, there is less evidence for the ER stress-dependent cluster formation at the endogenous levels of IRE1α (Sundaram et al., 2017).

Blue native poly acrylamide gel electrophoresis (BN-PAGE) immunoblotting has been successfully used to monitor the complex or oligomer formation of mitochondrial protein import machinery (Wittig et al., 2006). Recent studies have used BN PAGE to follow the dynamics of the Sec61 translocon complexes during the translocation into the ER lumen (Conti et al., 2011) and ligand-dependent oligomerization of NLRC4 inflammasome (Kofoed and Vance 2011). We have recently used this approach to specifically monitor the role of the Sec61 translocon in controlling IRE1α complexes (Sundaram et al., 2017). In the current study, we have employed this technique to investigate oligomerization dynamics of all three endogenous UPR sensors during ER stress. We found that all three UPR sensors existed as oligomeric complexes even under homeostatic conditions. BN-PAGE can robustly detect ER stress dependent changes in the oligomeric complexes of PERK and ATF6α. While the endogenous oligomeric complexes of IRE1α were not significantly changed during ER stress, a slight overexpression of IRE1α exhibited oligomerization changes in an ER stress dependent manner. Surprisingly, depletion of BiP had less impact on the oligomeric complexes of UPR sensors. Also, overexpressing BiP did not affect the oligomeric complexes of UPR, but significantly reduced all three UPR sensors sensitivity to respond to the accumulation of misfolded proteins. Thus, our results find less evidence for the BiP-mediated activation of UPR sensors, but rather support that misfolded proteins binding to preformed oligomers of UPR sensors may be crucial for activation.

## Results

### Changes in the endogenous complexes of UPR sensors under homeostatic and ER stress conditions

To monitor the changes in the endogenous complexes of IRE1α, PERK or ATF6α during homeostatic and ER stress conditions, we used a BN-PAGE immunoblotting procedure (Wittig et al., 2006; Sundaram et al., 2017). HEK293 cells were treated with thapsigargin (TG), which induces ER stress by inhibiting calcium transport into the ER, and prepared digitonin lysates for BN-PAGE immunoblotting. The activation of the endogenous IRE1α was monitored by probing its phosphorylation status using a phos-tag based immunoblotting (Yang et al., 2009; Sundaram et al., 2017). A significant proportion of IRE1α was activated after one hour of ER stress and inactivated within six hours of ER stress treatment (Figure 1A). In accordance with previous studies (Lin et al., 2007; Sundaram et al., 2017), PERK was activated throughout the stress period as shown by its phosphorylation (Figure 1A). ATF6α was activated upon ER stress as shown by the loss of signal due to the proteolytic release of the N-terminal fragment after its migration to the Golgi apparatus (Figure 1A). Similar to IRE1α, ATF6α was robustly attenuated within eight hours of stress period since the full-length ATF6α signal appeared back during the later hours of ER stress (Figure 1A).

**Figure 1.**
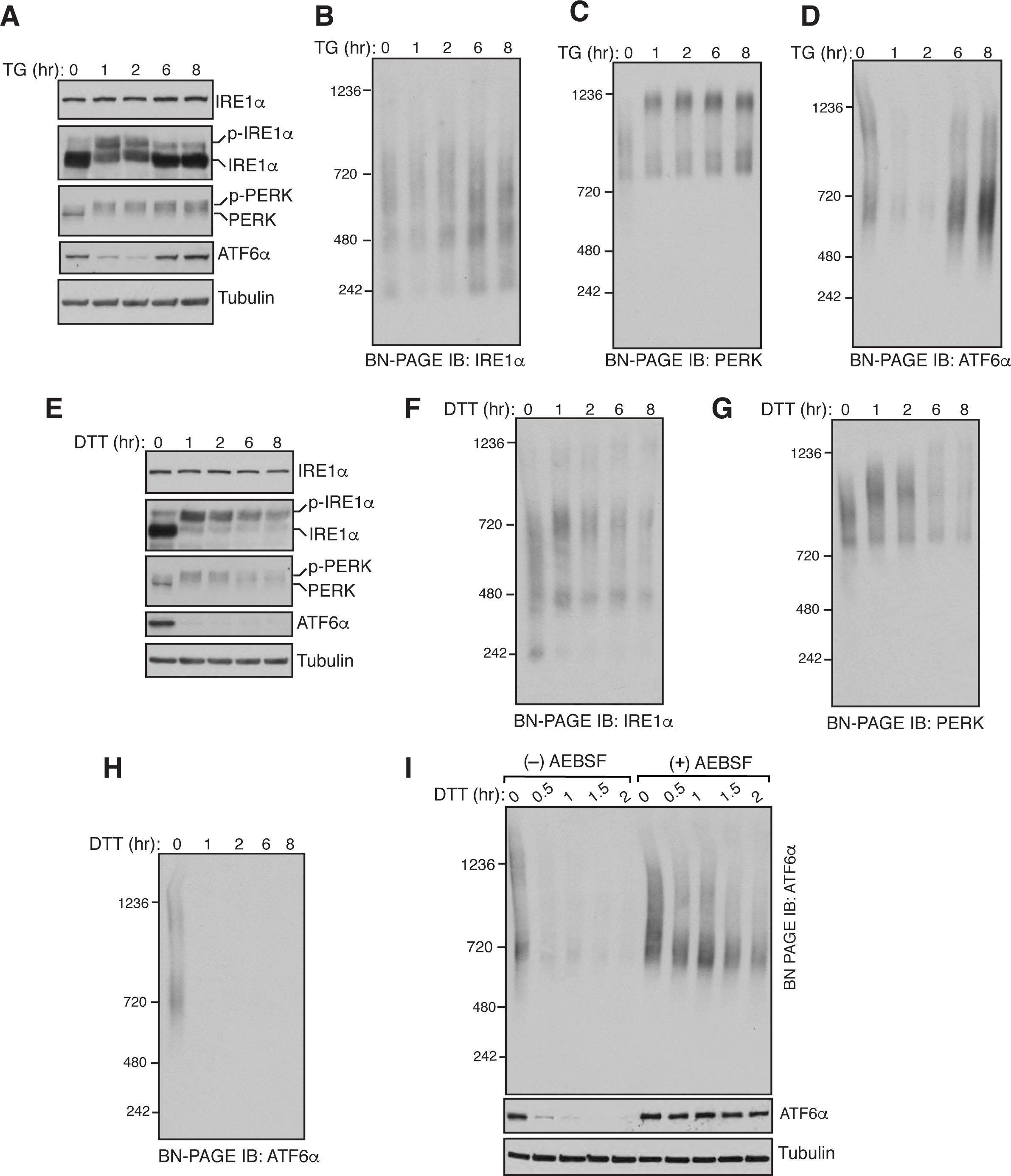
Changes in the endogenous complexes of IRE1α, PERK and ATF6α during ER stress. (A) HEK293 cells were treated with 7.5μg of thapsigargin (TG) for the indicated time points and lysed with digitonin. The lysates were separated by SDS PAGE and immunoblotting for the indicated proteins. A phos-tag based immunoblotting was performed for probing phosphorylated IRE1αα. (B) The digitonin lysates from A were analyzed by BN-PAGE immunoblotting with IRE1α antibodies, (C) PERK antibodies, or (D) ATF6α antibodies. (E-H) HEK293 cells were treated with 5mM DTT for the indicated time points and analyzed as in (A-D). (I) HEK293 cells were pretreated with a serine protease inhibitor AEBSF (250μM) for 1hr and subsequently treated with DTT at the indicated time to induce ER stress. The digitonin lysates were analyzed as in A and D. Experiments shown are representative of experiments repeated at least two times during different days.

Consistent with our previous findings (Sundaram et al., 2017), BN-PAGE immunoblotting revealed that IRE1α existed as predominantly two complexes: ~480 kDa and ~720 kDa complexes. Changes in IRE1α complexes were not apparent between unstressed and stressed cells, albeit the ~240kDa band become disappeared upon ER stress and appeared back in the later hours of ER stress (Figure 1B). ER stress dependent changes in PERK complexes were evident since PERK moved from a ~900 kDa complex to a ~1200 kDa complex upon ER stress (Figure 1C). BN-PAGE detected two large complexes of ATF6α under homeostatic conditions: ~720 kDa and ~1200 kDa, which were nearly disappeared during initial hours of ER stress and appeared back in the later hours of ER stress (Figure 1D). To rule out the possibility that oligomerization changes of UPR sensors on BN-PAGE are specific to TG treatment, we performed BN-PAGE analysis with cells treated with DTT, which induces protein misfolding in the ER by blocking protein disulfide bond formation. All three UPR sensors were robustly activated in cells treated with DTT (Figure 1E). In line with TG treatment, there were no significant changes in IRE1α complexes between unstressed and stressed cells (Figure 1F). Interestingly changes in PERK complexes were less noticeable with DTT treatment compared to TG treatment since not all the 900 kDa complex moved to the 1200 kDa complex (compare, Figure 1C and Figure 1G). ATF6α signal was disappeared throughout DTT treatment, suggesting that the ER is experiencing continuous stress and is not restored (Figure 1F).

Since ATF6α is proteolytically cleaved during ER stress, we were not able to detect the changes in ATF6α complexes. Furthermore, our ATF6α antibody failed to detect cleaved both N‐ and C-terminal fragments of ATF6α. To determine the changes in ATF6α complexes during ER stress, we inhibited S1P and S2P proteases that are responsible for the cleavage of ATF6α using a previously described serine protease inhibitor, 4-(2-aminoethyl) benzene sulfonyl fluoride hydrochloride (AEBSF) (Okada et al., 2003). In the presence of the inhibitor, ATF6α cleavage was nearly abolished as shown by immunoblotting (Figure 1I, bottom). Interestingly, the larger complex of 1200 kDa band significantly reduced during ER stress, whereas little change occurred with the smaller complex of 720 kDa (Figure 1I), suggesting that ER stress dependent decreased oligomerization of ATF6α is necessary for its transport to the Golgi apparatus.

Since all three UPR sensors appeared as large complexes on BN-PAGE, we wanted to exclude the possibility that the slow migration of UPR sensors on BN-PAGE was caused by their association with lipid or/and detergent micelles. We therefore tested the endogenous complexes of UPR sensors by a chemical crosslinking approach. HEK293 cells were treated with a cysteine reactive crosslinker and analyzed by a low percentage standard SDS PAGE immunoblotting. Remarkably, consistent with BN-PAGE data, all three UPR sensors entirely shifted to high molecular weight crosslinked adducts both in unstressed and stressed cells (Figure 1-figure supplement A, B, and C). Although BiP is a vastly abundant chaperone than all three UPR sensors, it showed significantly less crosslinked adducts compared to UPR sensors (Figure 1-figure supplement 1D). Given that the total protein profile did not significantly change with all concentrations of the crosslinker suggests that only stable oligomers like UPR sensors can be efficiently crosslinked at these concentrations (Figure 1-figure supplement 1E). For IRE1α and ATF6α, we do not expect to detect changes in crosslinked adducts between unstressed and stressed cells because the former did not change with stress on BN-PAGE, and the signal for the later mostly disappeared with stress. Interestingly, PERK also did not show any noticeable change in crosslinked adducts between unstressed and stressed cells (Figure 1-figure supplement 1B). The precise reason for this is unclear, but it is likely due to the limited resolution of the SDS PAGE to differentiate ~900 kDa complex of PERK in unstressed cells from ~1200 kDa complex in stressed cells. Collectively, these results suggest that all three UPR sensors existed as preformed oligomeric complexes and become activated upon ER stress by changing the oligomerization status for both PERK and ATF6α, but little to no change in IRE1α oligomerization at the endogenous levels.

### Overexpression of IRE1α exhibits ER stress dependent changes in its complexes

Although ER stress dependent changes in UPR complexes were obvious for both PERK and ATF6α, we were not able to detect changes in IRE1α complexes. This was surprising to us since previous studies have suggested that stress dependent higher order oligomerization is important for IRE1α activity (Li et al., 2010). We therefore increased the intensity of ER stress to detect changes in IRE1α complexes. All three UPR sensors were robustly activated from low to high concentrations of DTT treatment (Figure 2A). Surprisingly, even at a high dosage of DTT treatment, we did not notice appreciable changes in IRE1α complexes (Figure 2A). However, the size of PERK complexes enhanced with increasing concentration of the stress, whereas ATF6α signal disappeared at all concentrations of DTT treatment (Figure 2, C and D).

**Figure 2.**
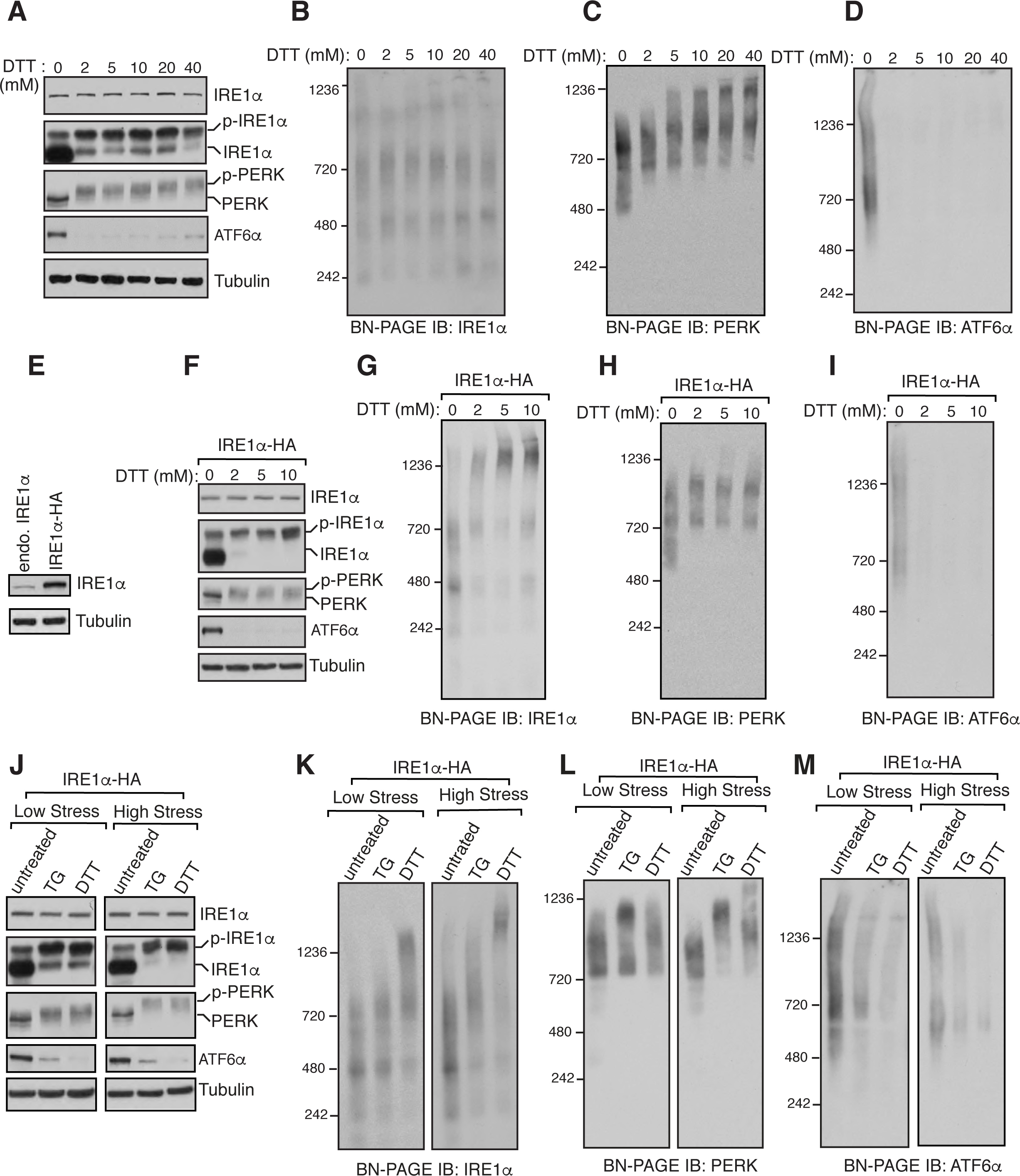
Overexpressed IRE1α, but not the endogenous IRE1α, exhibits ER stress induced changes in its complexes. (A) HEK293 cells were treated with the indicated concentrations of DTT for 2.5 hours and analyzed for immunoblotting with the indicated antigens. (B-D) The samples from A were analyzed by BN-PAGE immunoblotting and probed for the indicated antigens. (E) An immunoblot comparing the expression levels between the endogenous IRE1α in HEK293 and IRE1α-HA complemented into HEK293 IRE1α‐/‐ cells. (F) HEK293 IRE1α‐/‐ complemented IRE1α-HA cells were treated with increasing concentration of DTT for 2.5 hrs and analyzed by immunoblotting for the indicated antigens. (G-I) The samples from F were analyzed by BN-PAGE immunoblotting and probed for the indicated antigens. (J) HEK293 IRE1α‐/‐ complemented IRE1α-HA cells were induced with either low stress by treating with a low concentration of TG (2.5μg/ml) or DTT (2mM) or high stress by treating with a high concentration of TG (25μg) or DTT (30mM) for 2.5 hours. The cell lysates were analyzed by standard immunoblotting for the indicated antigens. (K-M) The samples from J were analyzed by BN-PAGE immunoblotting for the indicated antigens. Most of the experiments shown are representative of experiments repeated at least two times during different days.

We hypothesized that ER stress dependent changes were not detected for IRE1α complexes since the concentration of the endogenous IRE1α is extremely low to form higher order oligomers (Kulak et al., 2014). To test this, we used HEK293 IRE1α‐/‐ cells complemented with recombinant IRE1α, expression of which is relatively higher than the endogenous IRE1α, but it showed only a little constitutive activation under homeostatic conditions (Figure 2, E and F). In supporting our hypothesis, the overexpressed IRE1α exhibited an ER stress dependent change in its complexes on BN-PAGE since a large 1200 kDa complex appeared with increasing concentrations of DTT (Figure 2G). By contrast to TG treatment, ER stress dependent change was less noticeable for PERK complexes, whereas ATF6α signal disappeared upon treating with DTT (Figure 2, H and I). We next simultaneously compared the effect of TG or DTT treatment that had on recombinant IRE1α complexes. At low and high-stress treatment with either TG or DTT resulted in efficient activation of all three UPR sensors (Figure 2J). Consistent with our previous studies (Sundaram et al., 2017), ER stress dependent changes in recombinant IRE1α complexes were not obvious under low-stress conditions with TG treatment, but a modest increase in the size of IRE1α complexes occurred under high-stress conditions with TG treatment (Figure 2K). However, ER stress dependent increase in the size of recombinant IRE1α complexes was conspicuous under both low and high-stress conditions with DTT treatment (Figure 2K). The size of PERK complexes increased under both under low and high-stress conditions with TG treatment, but their size increase was apparent with only high-stress conditions with DTT treatment (Figure 2L). As expected, ATF6α complexes were responsive to both low and high-stress conditions as ATF6α signal disappeared under both conditions (Figure 2M). Together these results suggest that the activation of endogenous IRE1α does not require a significant change in its oligomeric complexes.

### Depletion of BiP has little effects on complexes and activation of UPR sensors

Since BN-PAGE can detect ER stress dependent changes in complexes of IRE1α, PERK, or ATF6α, we wanted to test the role of BiP in regulating oligomerization of these complexes in cells. Since BiP has been suggested to be a negative regulator by inhibiting the oligomerization and activation of UPR sensors (Okamura et al., 2000; Bertolotti et al., 2000), we expected that depletion of BiP might lead to significant changes in oligomeric complexes of UPR sensors in both unstressed and stressed cells. On the other hand, the depletion of Sec61 translocon would serve as a control since its depletion selectively changes IRE1α complexes as well as activates IRE1α (Sundaram et al., 2017). We therefore transiently depleted BiP or the Sec61 translocon in cells using siRNA-mediated knockdown. Probing the phosphorylation status of IRE1α and PERK revealed that a small amount of IRE1α and PERK were activated in BiP depleted cells during unstressed conditions and that became fully activated upon ER stress (Figure 3A). Depletion of BiP appeared to have little to no effect on the cleavage of ATF6α in unstressed as well as stressed cells relative to control siRNA depleted cells (Figure 3A). Consistent with the previous studies (Adamson et al., 2016), depletion of the Sec61 translocon selectively activated about 50% of IRE1α in unstressed cells, and that became fully activated upon ER stress. While the depletion of the Sec61 translocon did not affect PERK, a significant loss of ATF6α signal occurred relative to the control (Figure 3A). Since our ATF6α antibodies only detect the uncleaved form of ATF6α, we were not able to validate whether the loss of signal represented the activation of ATF6α or the level of ATF6α was reduced owing to the depletion of the Sec61. ATF6α was also not efficiently activated in the Sec61 translocon depleted cells upon ER stress since it remains predominantly uncleaved upon ER stress (Figure 3A). Intriguingly, the levels of IRE1α and PERK were slightly increased upon either depleting BiP or the Sec61 translocon in cells (Figure 3A).

**Figure 3.**
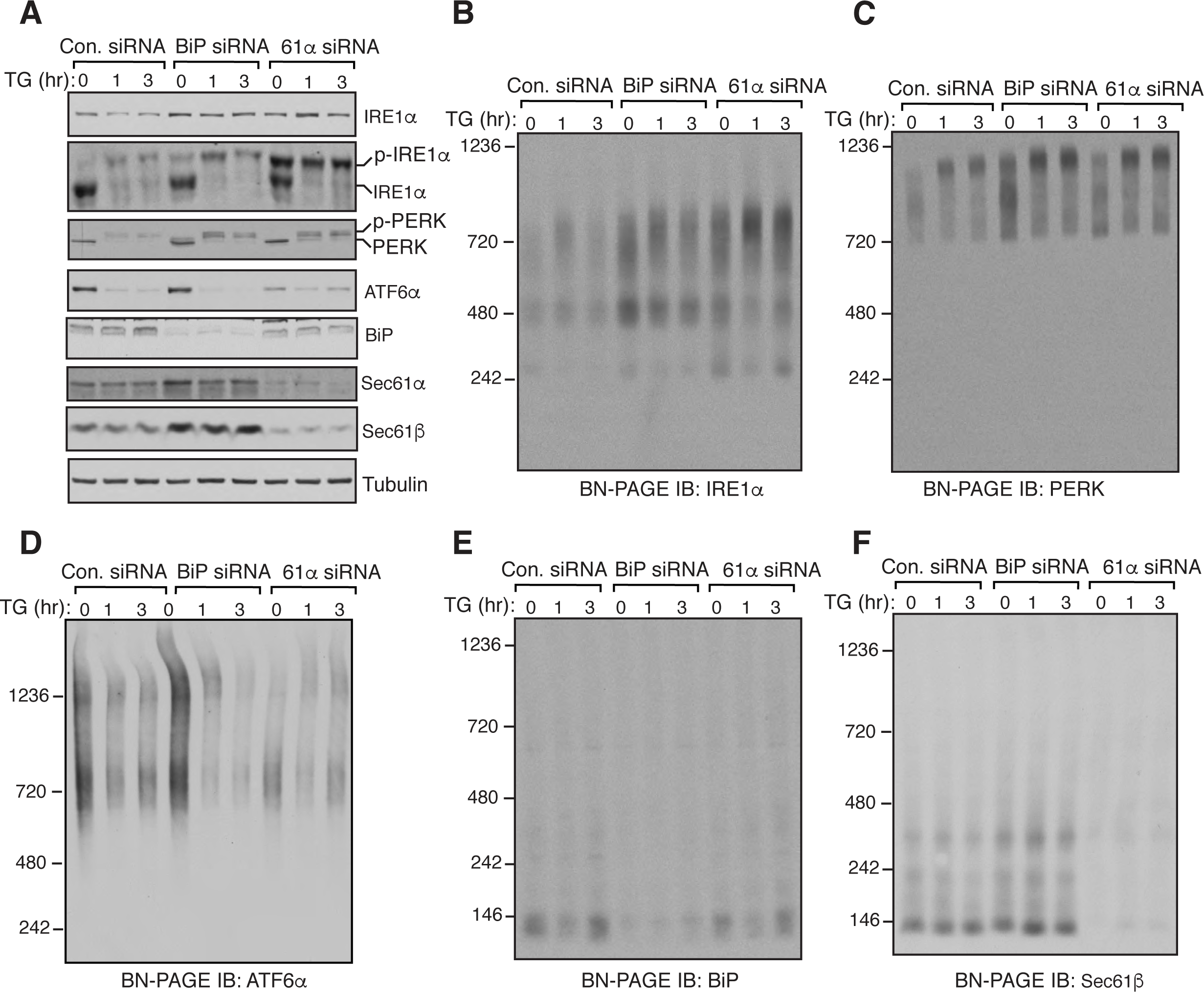
Depletion of BiP neither affects complexes or activation of all three UPR sensors. (A) HEK293 cells were transfected with siRNA oligos directed against either BiP siRNA or Sec61α for two consecutive days. After 72 hours of transfection, the cells were treated with 2.5μg of TG for the indicated time points and analyzed by immunoblotting for the indicated antigens. (B-F) The samples from A were analyzed by BN-PAGE immunoblotting for the indicated antigens. Experiments shown are representative of experiments repeated at least two times during different days.

BN-PAGE analysis of BiP depleted cells revealed no significant changes occurred with complexes of all three UPR sensors in both unstressed cells and stressed cells in comparison to control siRNA treated cells (Figure 3, B-D). Consistent with our recent studies (Sundaram et al., 2017), depletion of the Sec61 translocon led to a enrichment of 720 kDa complex of IRE1α compared to control or BiP siRNA treated cells both under normal and stress conditions (Figure 3B). In contrast, the Sec61 translocon depletion did not affect either PERK or ATF6α complexes (Figure 3C, D). Unlike all three UPR sensors, BiP migrated as predominantly a single species ~140 kDa on BN-PAGE, whereas the Sec61 translocon ran predominantly as a ~140 kDa form as well as a ~ 350 kDa form, which is in agreement with previous studies (Figure 3E, F) (Conti et al., 2015; Sundaram et al., 2017). Together these data suggest that the depletion of BiP had minor effects on the complexes and activation of all three UPR sensors both under unstressed and stressed conditions.

### BiP overexpression does not impact complexes of UPR sensors but suppresses activation of UPR sensors

We next tested whether overexpression of BiP would reduce the size of UPR complexes as well as blocks the activation of UPR sensors. Consistent with the previous studies (Kohno et al., 1993; Bertolotti et al., 2000), overexpression of BiP above the endogenous level significantly suppressed the activation of IRE1α as reflected by a significantly reduced phosphorylation of IRE1α upon TG induced ER stress (Figure 4A). To our surprise, PERK was activated as shown by its phosphorylation status even in BiP overexpressing cells treated with TG, albeit slightly less efficient than the control. Interestingly, the ATF6α protein level was about two-fold increased in BiP overexpressing cells and that the ER stress dependent cleavage of ATF6α was nearly blocked (Figure 4A). BN PAGE analysis of IRE1α in BiP overexpressing cells revealed that the overall pattern of IRE1α complexes was not significantly affected in the presence or absence of the TG treatment (Figure 4B). However, we noticed a large complex (~1200 kDa) that specifically formed for IRE1α upon BiP overexpression in a non-ER stress dependent manner. BiP overexpression also did not affect the size of PERK complexes in unstressed cells. Consistent with the activation data, PERK normally moved from a smaller complex to a large complex on BN-PAGE in BiP overexpressing cells upon treatment with TG (Figure 4, A and B). ATF6α complexes were not affected by BiP overexpression in unstressed cells, but it blocked the disappearance of ATF6α complexes upon ER stress (Figure 4D). These results were further corroborated by BiP overexpressing cells treated with DTT (Figure 4, E-H). Nevertheless, we observed one particular difference for PERK between TG and DTT treated cells. While the activation of PERK was not significantly affected by BiP overexpression in TG treated cells, it nearly blocked activation of PERK upon DTT treatment (Figure 4, E and G). Taken together, our data suggest that the overexpression of BiP does not impact the oligomeric complexes of all three UPR sensors, but interferes with the activation of UPR sensors upon ER stress, presumably by sequestering misfolded proteins away from UPR sensors.

**Figure 4.**
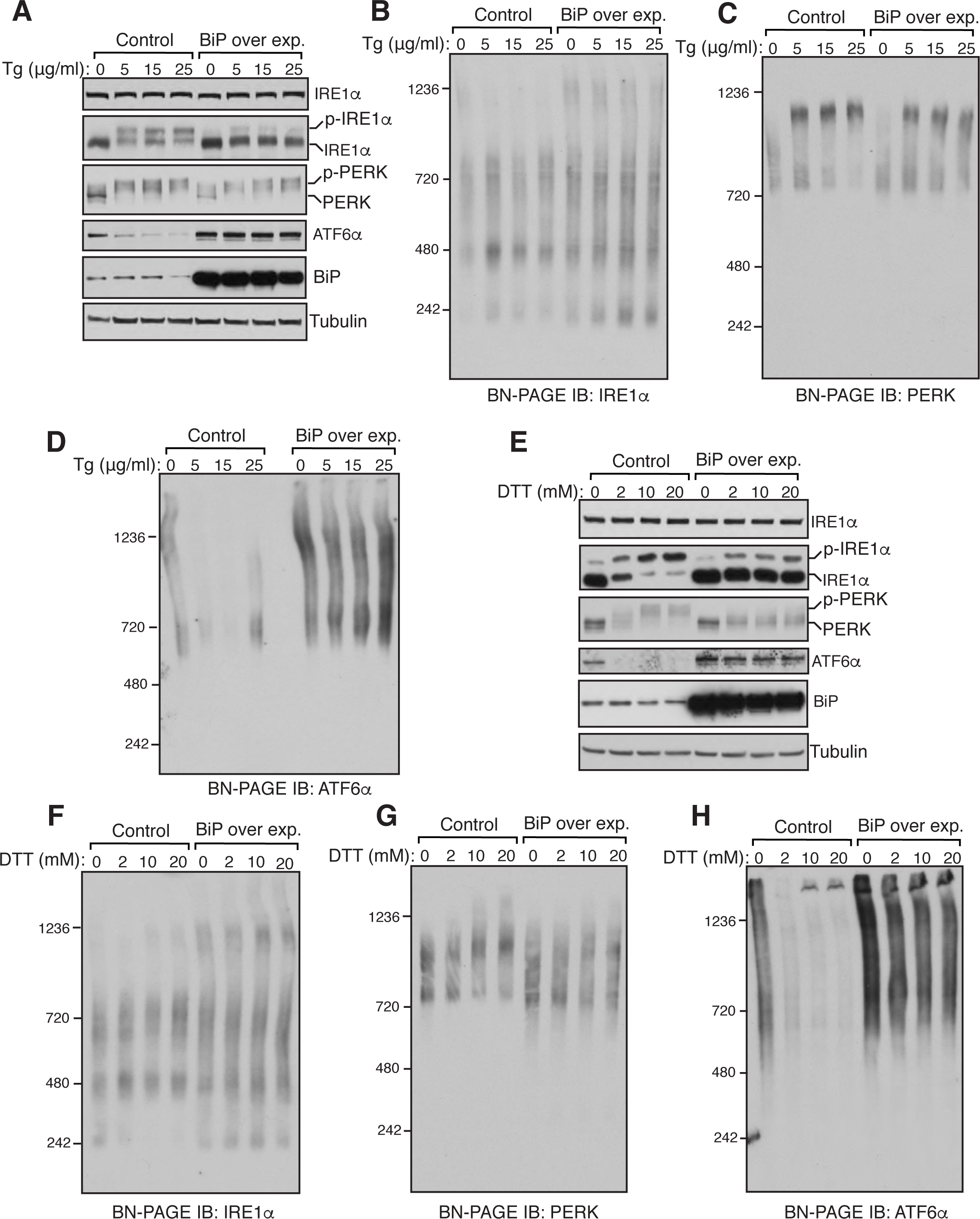
Overexpression of BiP has little effects on complexes of IRE1α, PERK, and ATF6α. (A) HEK293 cells were transfected with either an empty plasmid (control) or plasmid encoding BiP. After 16 hours of transfection, media was replenished and grown for another 24 hrs. The cells were treated with the indicated concentrations of TG for 2 hrs. The cells were harvested and analyzed by immunoblotting for the indicated antigens. (B-D) The samples from A were analyzed by BN-PAGE immunoblotting for the indicated proteins. (E) As in A, the cells were transfected with either an empty plasmid or plasmid encoding BiP and treated with the indicated concentrations of DTT for 2 hrs. The cells were harvested and analyzed by immunoblotting for the indicated antigens. (F-H) The samples from E were analyzed by BN-PAGE immunoblotting for the indicated proteins. Experiments shown are representative of experiments repeated at least two times during different days.

### IRE1α can interact with misfolded proteins in cells

The results above argue against the model that BiP binds to monomers of UPR sensors and inhibits constitutive oligomerization and activation. We therefore favour the model that misfolded proteins might bind and activate the preformed oligomeric complexes of UPR sensors, which would have a higher affinity for misfolded proteins. To support this model, we attempted to capture the interaction between the UPR sensors and misfolded proteins in the ER lumen. However, we failed to see the interaction by coimmuoprecipitation experiments. We reasoned that either the interaction is weak or it is sensitive to immunoprecipitation conditions. To circumvent this issue, we first expressed a misfolded alpha1-antitrypsin variant, null (Hong Kong) (NHK), into HEK293 IRE1α‐/‐ complemented with HA-tagged IRE1α and induced ER stress with DTT. The cells were then treated with a reversible lysine reactive crosslinker. The crosslinked samples were immunoprecipitated for IRE1α using an HA antibody. Indeed, we noticed an interaction between IRE1α and misfolded antitrypsin (Figure 5A), which was slightly improved with ER stress. As previously reported (Bertolotti et al., 2000; Oikawa et al., 2009), we noticed an interaction between IRE1α and BiP, which was reduced as the intensity of the stress increased (Figure 5A). We obtained a similar result when we treated cells with TG (Figure 5B). Thus, the interaction between misfolded antitrypsin and IRE1α suggest that misfolded proteins bind and activate IRE1α and likely other UPR sensors.

**Figure 5.**
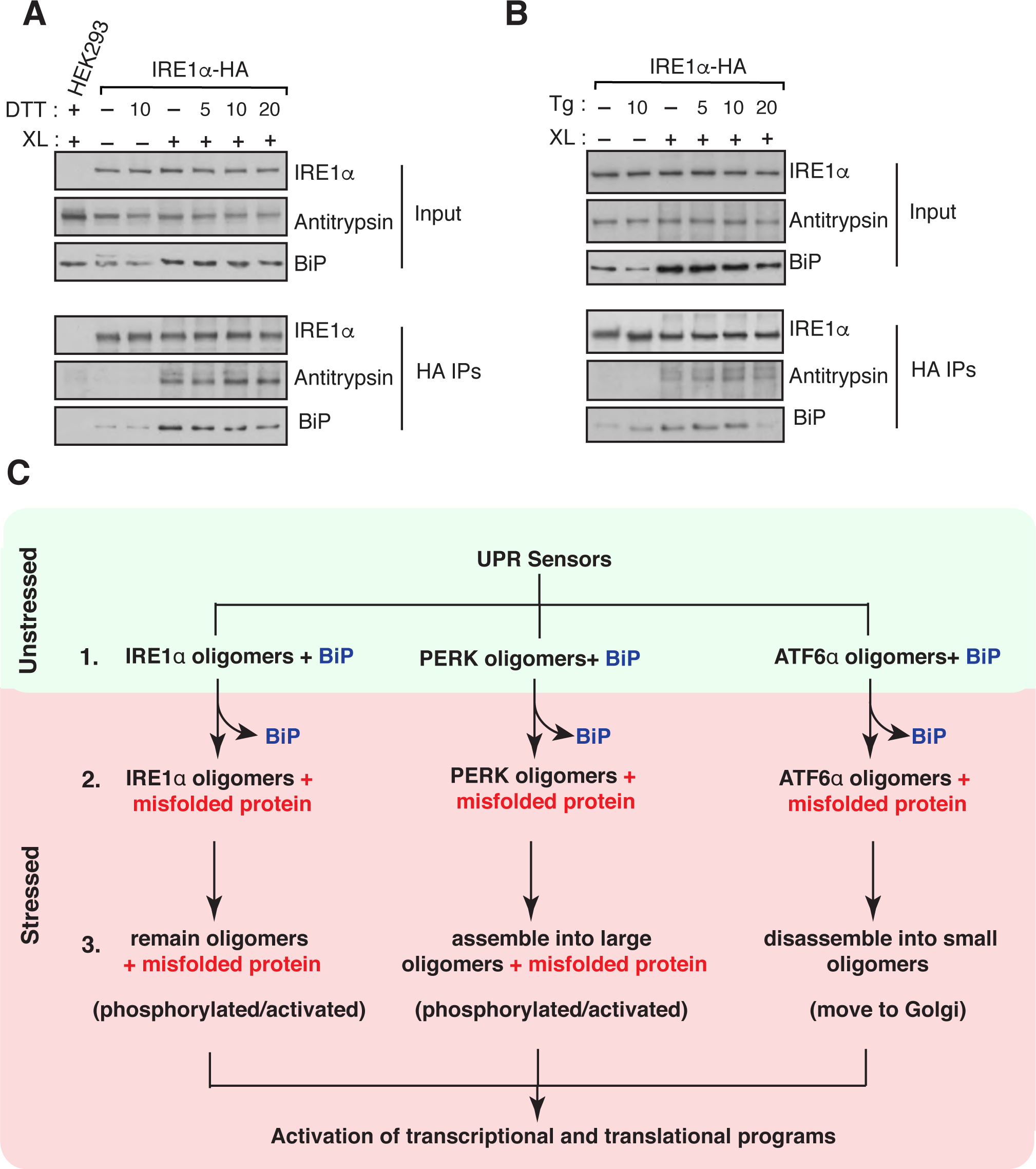
The UPR sensor IRE1α interacts with misfolded antitrypsin. (A) HEK293 IRE1α‐/‐ cells complemented with IRE1α-HA were transfected with either empty plasmid or NHK a1 antitrypsin and induced with 10 ng of doxycycline to drive the expression of IRE1α-HA. After 24 hrs of transfection, the cells were reacted with DSP crosslinker (XL). As a control, HEK293 cells transfected NHK α1 antitrypsin and treated with the crosslinker. The crosslinked samples were immunoprecipitated with anti-HA magnetic antibodies and analyzed by immunoblotting with the indicated antibodies. (B) The procedure was identical to A, but the cells were treated with 5μg TG for 1 hr before crosslinking. Experiments shown are representative of experiments repeated at least two times during different days. (C) Models for oligomerization and activation of UPR sensors. Step 1: All three endogenous UPR sensors exist as preformed oligomers associated with BiP in cells. Step 2: Upon ER stress, oligomers of UPR sensors bind to misfolded proteins with concomitant release of BiP from UPR sensors. Step 3: Once binding to misfolded proteins, IRE1α may undergo conformational changes without major changes in the oligomerization state, which in turn activates its kinase and RNase activities. PERK is activated and phosphorylated through assembling into large oligomers from small oligomers upon binding with misfolded proteins. Conversely, misfolded proteins binding to ATF6α oligomers induce disassembly of oligomers, thus migrating to Golgi for activation.

## Discussion

In mammalian cells, three UPR branches, IRE1α, PERK and ATF6α become activated upon accumulation of misfolded proteins in the ER (Walter and Ron, 2011). Once activated, these UPR sensors initiate transcriptional and translational programs to improve the protein folding capacity of the ER. How these UPR sensors detect the accumulation of misfolded proteins, and how they become activated have been debated in the field (Snapp, 2012). In the first model, the luminal sensor domains of all three UPR sensors bind to BiP under homeostatic conditions (Bertolotti et al., 2000; Okamura et al., 2000; Shen et al., 2002; Carrara et al., 2005). As misfolded proteins accumulate during ER stress, BiP releases, and the UPR sensors become activated. In the second model, misfolded proteins directly bind and activate all three UPR sensors during ER stress (Gardner and Walter, 2011; Gardner et al., 2013). In both models, oligomerization changes in UPR sensors appear to play a crucial role in activation. One critical barrier to test these different models is to reliably monitor the changes in oligomerization status of all three UPR sensors during ER stress conditions. In the present study, we have monitored all three UPR sensors side by side by employing a BN-PAGE immunoblotting technique. This method allows us to directly compare changes in complexes of all three UPR sensors in unstressed and stressed cells (Figure 5C).

Our BN-PAGE data suggest that all three UPR sensors existed as preformed oligomers under homeostatic conditions. There are two alternative possibilities to this claim. First, it is likely that preformed oligomers of UPR sensors may reflect their interaction with their partner proteins in cells. Second, it is possible that the UPR sensors migrate slower in the BN-PAGE due to their association with lipids and detergent micelles, although all three sensors have only a single transmembrane domain. However, there are several of our observations argue against these alternative possibilities. First, our crosslinking data provide evidence that all three endogenous UPR sensors completely shifted to larger size crosslinked adducts upon chemical crosslinking, suggesting that they form stable oligomeric complexes. Second, BiP is a known interacting protein of all three UPR sensors, but its depletion does not result in any apparent changes in the size of UPR complexes, suggesting that interacting proteins of UPR sensors may not significantly impede their migration on BN-PAGE. Third, the fact that IRE1α and to a lesser extent PERK with DTT treatment can be activated with no significant changes in oligomerization suggest that they are already in preformed oligomers since the monomeric form of IRE1α is inactive (Zhou et al., 2006; Li et al., 2010). Furthermore, our BN-PAGE data with ATF6α is consistent with previous studies that the ATF6α appears to be in higher order oligomers under homeostatic conditions (Nadanaka et al. 2007; Gallagher et al., 2016).

The endogenous IRE1α complexes exhibit little changes upon activation by ER stress treatment even at high concentrations of DTT. This data is contrary to the current model that IRE1α is proposed to be in monomers in unstressed cells and becomes oligomerized for activation upon ER stress (Kimata and Kohno, 2011; Hetz, 2012; Walter and Ron, 2011).It is unlikely that BN-PAGE cannot detect IRE1α oligomerization status since it can apparently detect the oligomerization changes for PERK and ATF6α. Moreover, ER stress dependent oligomerization changes for IRE1α can be observed with a slight overexpression of IRE1α. At present, it is unclear why the endogenous IRE1α cannot be further oligomerized upon ER stress. One possibility is that there are not sufficient oligomers of IRE1α in the ER membrane to form higher order oligomers upon ER stress. Alternatively, unlike PERK and ATF6α, the Sec61 translocon represses IRE1α oligomerization, thereby attenuating IRE1α activity during ER stress (Plumb et al., 2015; Sundaram et al., 2017). However, overexpression of IRE1α can form ER stress dependent higher order oligomers since the overexpression results in free IRE1α oligomers that are not associated with Sec61 (Plumb et al., 2015). Interestingly, ER stress dependent changes in oligomerization of recombinant IRE1α are apparent with DTT treatment, but less noticeable with TG treatment, suggesting that IRE1α is more responsive to DTT treatment. While PERK is robustly oligomerized and activated by TG induced ER stress, it is slightly less sensitive to DTT induced oligomerization and activation, which is in agreement with the earlier studies (DuRose et al., 2006).

ATF6α signal quickly returns within six hours of ER stress treatment, suggesting that the activation of ATF6α is attenuated despite the presence of ER stress. In contrast to previous findings (Lin et al., 2007), ATF6α inactivation rate is very similar to attenuation of IRE1α signalling during ER stress. This discrepancy may be due to the use of heterologously expressed ATF6α in the previous studies, which responds inefficiently to ER stress, whereas the endogenous ATF6α in the current study and studies from the Mori group show a robust activation and inactivation to ER stress treatments (Okada et al., 2003). Unlike IRE1α and PERK, changes in oligomerization of ATF6α cannot easily be monitored since the loss of ATF6α signal owing to the release of the cytosolic N-terminal domain of ATF6α from its membrane anchor domain during ER stress. However, the inhibition of ATF6α cleavage by AEBSF (Okada et al., 2003) proves to be a useful tool to monitor ER stress dependent conversion of two different ATF6α oligomeric complexes to a single oligomeric complex. It remains to be determined whether the protease inhibitor has any secondary effects on the oligomerization of ATF6α.

Our ability to monitor stress dependent changes in all three UPR sensors in parallel motivated us to test the role of BiP in inhibiting the oligomerization of these sensor proteins. According to this model, we predicted that the depletion of BiP should enhance the oligomerization of UPR sensors, whereas overexpression of BiP should reduce the size of UPR oligomers even under normal conditions. However, we found no significant changes in oligomerization of all three UPR sensors upon significant depletion of BiP in cells. This is also further supported by our observation that all three UPR sensors are largely remain inactive upon BiP depletion. Furthermore, it is unclear why the depletion of BiP leads to slightly increased protein levels for IRE1α and PERK. It is likely that BiP may be involved in the turnover of these UPR sensors since a recent study implicates BiP in the degradation of IRElα (Sun et al., 2015). It remains to be seen whether the residual amount of BiP in the depleted cells is sufficient to bind and suppress the activation of UPR sensors under homeostatic conditions.

Consistent with the previous studies, the overexpression of BiP effectively suppressed all three UPR sensors under DTT stress conditions (Kohno et al., 1993; Bertolotti et al., 2000). Surprisingly, PERK can still be activated by TG treatment, whereas the activation of IRElα and ATF6α are suppressed. This result implies that PERK is the most sensitive arm of the UPR and such that it is also not easily attenuated during ER stress conditions. Furthermore, the overexpression BiP has no impact on reducing the size of oligomers of UPR sensors. Taken together, these results raise the question, how are UPR sensors activated? Our data point to the peptide-binding model (Credle et al., 2005), where UPR sensors are activated by directly binding to misfolded proteins (Figure 5C). Importantly, our findings of preformed oligomers of UPR sensors in unstressed cells may explain the robust nature of UPR response even at low levels of ER stress (Rutkowski et al., 2006) since oligomers of UPR sensors would have a higher affinity for misfolded proteins (Gardner et al., 2011). Our crosslinking data that captures the interaction between IRE1α and a misfolded protein further support the peptide-binding model. However, it remains to be determined whether PERK and ATF6α can also interact with misfolded proteins. In this model, the interaction between UPR sensors and BiP might play an important role by increasing the local BiP concentration around UPR sensors, thus preventing inappropriate activation during homeostatic conditions. Future studies should focus on how binding of misfolded proteins could activate UPR sensors using assays that use physiological concentration of purified full-length UPR sensor proteins. It is also necessary to consider using different misfolded substrates for binding studies with UPR sensors since each UPR sensor may prefer different misfolded substrates.

## Materials and methods

### Antibodies and Reagents

Antibodies were purchased: anti-IRE1α (3294, Cell Signaling, RRID:AB_823545), anti-PERK (3192, Cell Signaling, RRID:AB_2095847), anti-IRE1α (20790, Santa Cruz, RRID:AB_2098712 ), anti-Tubulin (ab7291, Abcam, RRID:AB_2241126), anti-BiP/GRP78 (610979, BD Biosciences, RRID:AB_398292). Anti-HA, anti-Sec61α, and anti-GFP were a gift from Dr. Ramanujan Hegde. Anti-mouse Goat HRP (11-035-003, Jackson Immunoreserach), anti-rabbit Goat HRP (111-035-003, Jackson Immunoreserach, RRID: AB_2313567), and anti-HA magnetic beads (88836, Fisher scientific).

Reagents were purchased: DMEM (10-013-CV, Corning), FBS (89510-186, VWR), Penicillin/Streptomycin (15140122, Gibco), Lipofectamine 2000 (11668019, Invitrogen), Doxycycline (631311, Clontech), AEBSF (A8456, Sigma), TG (BML-PE180-0005, Enzo Life Sciences), Tunicamycin (T7765, Sigma), BMH (bismaleimidohexane) (22330, ThermoFisher), DSP (dithiobis(succinimidyl propionate)) (22585, ThermoFisher) Protease inhibitor cocktail (11873580001, Roche), Digitonin (300410, EMD Millipore), Phos-tag (300-93523, Wako), 3-12% BN-PAGE Novex Bis-Tris Gel (BN1003BOX, Invitrogen), SuperSignal West Pico or Femto Substrate (34080 or 34095, Thermo Scientific). All other common reagents were purchased as indicated in the method section.

### Cell culture and ER stress treatment

HEK 293-Flp-In T-Rex cells were cultured in high glucose DMEM (Corning) containing 10% FBS (Gibco), 100 U/ml penicillin and 100 μg/ml streptomycin (Gibco) at 5% CO2. IRE1αα‐/‐ HEK293-Flp-In T-Rex cells and IRE1α complemented cells were previously described (Plumb et al., 2015). The cells were transfected with plasmid or siRNA oligos using Lipofectamine 2000 according to manufacturer's protocol. For ER stress treatment, HEK293 cells were counted and plated in 6 well plates (1.5 × 10^6^) and grown overnight. The cells were then treated with either DTT or TG. All the concentrations and treatment time were as indicated in either result or figure sections.

For the depletion experiments, 0.5 × 10^6^ cells were plated and transfected next day with either BiP siRNA (GGAGCGCAUUGAUACUAGA) or Sec61α siRNA (Plumb et al., 2015). After 24 hours of first transfections, cells were again transfected with siRNA oligos. After 72 hours of the first transfection, cells were treated with TG as indicated in the figure legends. For BiP expression experiment, 0.5 × 10^6^ cells were plated and grown overnight. The cells were then transfected with transfected with rat BiP plasmid (a kind gift from Dr.Ramanujan Hegde) or empty vector. After 24 hours of transfection, cells were treated with ER stress inducers as mentioned in the figure legends. After the treatment, cells were washed with 1xPBS, harvested in 1.2ml 1xPBS. The cells were spun at 10,000g for 1 minute, and the pellets were flash frozen and stored at -80C.

### Chemical crosslinking

HEK293 cells were plated on six well plates (0.75 × 10^6^ cells/ well) and grown overnight. The cells were either left untreated or treated with 5μg TG/ml for 60 minutes. After the treatment, the cells were washed once with KHM buffer (20mM HEPES pH 7.4, 110mM NaCl, 2mM MgAc) and permeabilized with 0.005% Digitonin for 5min on ice. The digitonin buffer was removed and washed once with KHM buffer. Subsequently, the permeabilized cells on plates were treated with 0.5 to 10μM BMH in KHM buffer for 30 minutes on ice. The cells were directly harvested in 2X SDS sample buffer, boiled, separated on 6% Tris-Glycine based gels and immunoblotted with the indicated antibodies in the Figure S1.

For DSP crosslinking, HEK293 IRE1α‐/‐ cells complemented with IRE1α-HA were plated on six well plates (0.75 × 10^6^ / well). The cells were then transfected with NHK α 1 antitrypsin plasmid (a Kind gift from Dr. Peter Cresswell) and induced with 10 ng of Doxycycline to drive the expression of IRE1α-HA. After 24 hours of transfection, the cells were washed with KHM and treated with 1mM DSP for 30 minutes at room temperature. The crosslinking reaction was then quenched with 0.1M Tris pH 8.0 for 15 min and harvested in RIPA buffer by incubating for 30 minutes in the cold room. The cell lysates were centrifuged at 18,500g for 15 min at 4°C. The supernatant was incubated with anti-HA magnetic beads for 2 hours at 4°C, washed, eluted with 2X SDS sample buffer and processed for immunoblotting with the indicated antigens in the figure.

### BN-PAGE Immunoblotting

The cell pellets were lysed using 2% digitonin buffer (50mM BisTris pH 7.2, 1x protease inhibitor cocktail [Roche], 100mM NaCl and 10% Glycerol) for 45 minutes. The cell lysates were then diluted to a final concentration of 1% digitonin and 50mM NaCl and centrifuged at 18,500g for 20 minutes at 4°C. The supernatant was collected and mixed with BN-PAGE sample buffer (Invitrogen) and 5% G520 (Sigma).

The samples were run using 3-12% BN-PAGE Novex Bis-Tris (Invitrogen) gel at 150 V for 1 hour with the dark blue buffer (50mM Tricine pH 7, 50mM BisTris pH 7 and 0.02% G250) at room temperature. The dark blue buffer was then exchanged with the light blue buffer (50mM Tricine pH 7, 50mM BisTris pH 7 and 0.002% G250) for 4 hours in the cold room.

To probe BiP, the gels were run for 1 hour with the dark blue buffer at room temperature and 3 hours with the light blue buffer in the cold room. After electrophoresis, the gel was gently shaken in 1x Tris-Glycine-SDS transfer buffer for 20 minutes to remove the residual blue dye. The transfer was performed using PVDF membrane (EMD Millipore) for 1 hour and 30 minutes at 85V. After transfer, the membrane was fixed with 4% acetic acid and followed with a standard immuno blotting procedure.

### Phostag assay

IRE1α phosphorylation was detected by previously described method (Yang et al., 2010). Briefly, 5% SDS PAGE gel was made containing 25μM Phos-tag (Wako). SDS-PAGE was run at 100 V for 2 hours and 40 minutes. The gel was transferred to nitrocellulose (Bio-Rad) and followed with immunoblotting.

## Acknowledgement

We thank Mariappan lab for useful discussions. We thank Sha Sun for providing comments on the manuscript. A.S. was supported by the Rudolph J. Anderson Postdoctoral Fellowship. This work is supported by the Yale School of Medicine start-up package and NIH 1R01GM11738601.

## Figure legends

**Figure 1-figure supplement 1:**
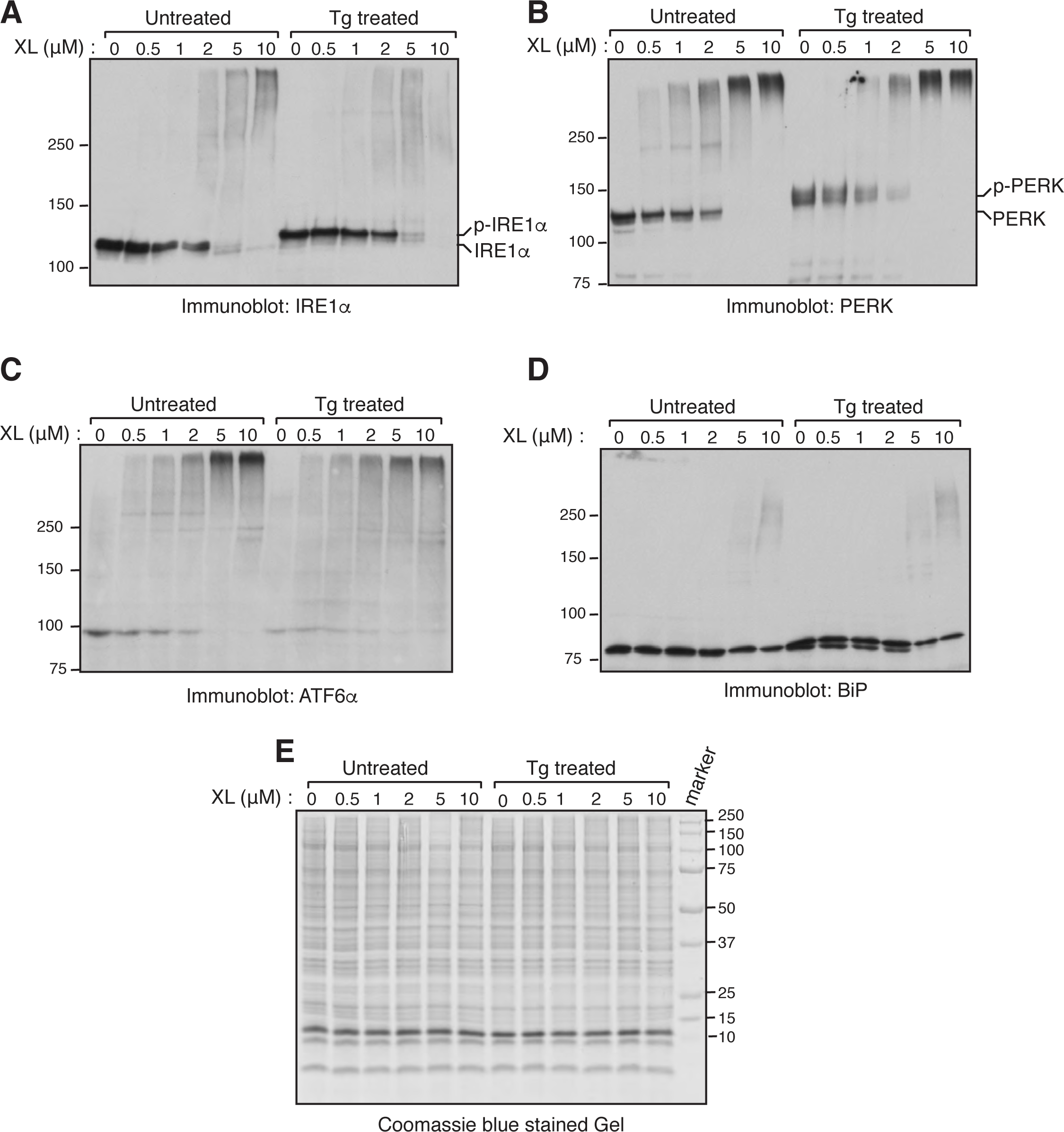
Chemical crosslinking of UPR sensors in cells. (A-E) HEK293 cells were either left untreated or treated with 5μg TG/ml for 60 min. After the treatment, the cells were permeabilized with a low concentration of Digitonin. Subsequently, the permeabilized cells were treated with 0.5 to 10μM BMH for 30 min on ice. The cells were directly harvested and analyzed by either immunobloting with the indicated antigens or coomassie blue staining.

